# A T-cell independent universal cellular therapy strategy through antigen depletion

**DOI:** 10.1101/2021.03.29.437606

**Authors:** Dan Li, Wenbing Wang, Shufeng Xie, Maolin Ge, Ruiheng Wang, Qiongyu Xu, Yan Sun, Jiang Zhu, Han Liu

**Affiliations:** Shanghai Institute of Hematology, State Key Laboratory of Medical Genomics, National Research Center for Translational Medicine at Shanghai, Rui Jin Hospital, School of Medicine and School of Life Sciences and Biotechnology, Shanghai Jiao Tong University, Shanghai, China

## Abstract

CD19-targeting chimeric antigen receptor (CAR) T-cell therapeutics is a revolutionary, novel and successful treatment for B-cell malignancies. However, while CD19-CAR-T therapy can obtain high rates of complete responses in these patients, a significant fraction of patients may experience CD19-negative relapse. Moreover, the dependency on T-cell mediated cytotoxicity restricts CAR-T therapy as a patient-specific individualized therapy with severe side effects such as cytokine-release syndrome (CRS). Whether CAR-T therapy can be substituted by a non-T-cell based universal cellular therapy is largely unknown. Surprisingly, we have demonstrated here that T-lymphocytic cells, as well as non-lymphocytic cells, can cause CD19 internalization and subsequent depletion when they are armed with a CD19-recognizing moiety. This CD19 antigen depletion can efficiently induce T-cell independent apoptosis in target cancer cells whose survival is dependent upon CD19 expression, suggesting that CD19 antigen depletion constitutes a crucial tumor destroying mechanism for CD19-CAR-T, especially for its long-term efficacy. We therefore proposed a universal strategy for CRS-free cellular therapeutics, utilizing artificial antigen-recognizing cells (AARC), which can be manufactured universally and standardly as “off-the-shelf” mesenchymal stromal cells (MSCs) or other types of non-autologous cell expressing anergic CARs. Our results not only uncovered an unrecognized mechanism for CAR-T cytotoxicity and antigen loss, but also shed new insight into a shift in cellular therapeutics from unique patient-specific autologous therapeutics, to universal and standardized allogeneic treatment.

## Introduction

Autologous CAR-T cells constitute a promising novel therapeutic approach for the treatment of hematological malignancies. Although this individual therapy-based approach has resulted in outstanding clinical results to date, it has several intrinsic disadvantages, such as the risk of manufacturing failure in certain patients, the risk of delay in treatment due to time costing manufacturing procedures, as well as financial burdens. As a result, the development of the next generation allogeneic cellular therapeutic strategy to address these issues is an active area of research.^1^

Cytotoxic T lymphocytes (CTL) mediate their anti-tumor effects predominantly through the granule exocytosis axis, the death ligands axis, as well as the release of cytokines.^2^ As one of the main adoptive T-cell therapy (ACT) approaches, CAR T cells also mediate target cell death mainly through these pathways.^3^ Most adoptive studies have focused on CD8+ T-cell populations because of their direct cytolytic activity. However, preclinical and clinical data have established the importance of incorporating CD4+ T-helper cells during immunotherapy.^4^ Moreover, adoptive transfer of CD4+ T-cell populations has shown that these cells can mediate tumor regression.^5^ Interestingly, although CD4+ CAR T cells are slower than CD8+ CAR T cells in terms of target cell death, single-cell analysis revealed equal amounts of tumor cell death was achieved by these cells.^6^ These results suggested that the direct cytolytic activity may not be absolutely required for CAR T-cell mediated target cell death. It has been proposed that CD4+ T-helper cells can aid the survival of infused CD8+ T cells, induce tumor cell senescence through derived cytokines, and even differentiate into cytolytic effectors.^7–9^ Nevertheless, whether T-cell independent mechanisms are involved in anti-tumor effects has not yet been addressed.

Successful CAR-T therapeutics depends heavily upon the in vivo activation and expansion of CAR-T-cell products, which inevitably cause the well-characterized side effect of cytokine release syndrome (CRS). Therefore, the development of T-cell independent universal cellular therapy strategies may not only provide an alternative option for “off-the-shelf” and standardized treatments, but also reduce the risk of CRS. Here, we have demonstrated that CAR-T cells can destroy target cells through a T-cell independent mechanism. Based on this finding, we proposed the concept of a universal cellular therapy strategy, which could serve as a companion to current CAR-T therapeutics.

## Methods

### Cell lines and primary cells

SEM, REH, RAJI, Jurkat, and K562 cell lines were obtained from DSMZ. Mesenchymal stem cells (MSCs) were obtained from Shanghai Nerostem Tech. CD3+ T cells were isolated using EasySep Human T Cell Isolation Kit (STEM CELL Technologies) and then cultured in CTS T Cell Expansion medium (Thermo) containing 10% FBS and 100 IU/ml human IL-2 (PeproTech). The CellTiter 96 MTS assay (Promega) was used to determine the cell viability and proliferation.

### Plasmid constructions

Fragments encoding CD19, CD22 and CD133-specific competent CARs and anergic CARs (scFvs) that lacks co-stimulatory and ζ-chain signaling domains were inserted into the lentiviral vector pCDH-T2A-copGFP (System Biosciences). The CD19– mRuby2 fusion was generated by fusing the mRuby2 sequence at the C terminus of CD19 and cloned into the pCDH lentiviral vector. Target sequences (CTTCAACGTCTCTCAACAGAT #1 and CCGAGTTCTATGAGAACGACT#2) against human CD19 and a control scrambled sequence (CTCAATCAACAGATCTCGTCT) were inserted into the pLKO.1 vector (Sigma).

### Flow cytometry

For cell labelling, the the Cell Trace Far Red Proliferation Kit and the Cell Trace CFSE Cell Proliferation Kit (Invitrogen) was used. Human CD19-APC and CD69-APC antibodies were obtained from BD Biosciences. Human CD133-PE antibody were purchased from Miltenyi Biotec. Human CD22 antibody were obtained from Biolegend. Apoptosis was measured using the Annexin V Apoptosis Detection Kit (BD Bioscience). Flow cytometry was performed on LSRFortessa or FACSAria sorter (BD Biosciences). Data were analyzed by the FlowJo software.

### Reagents

Bortezomib (Velcade), Sc-79, CsA, and dynago-4a were obtained from Selleck Chemicals. Bafilomycin A1 (Baf-A1), DC661, MK-2206, and MβCD were obtained from MedChemExpress.

### Immunoblots

Human CD19 and Akt antibodies were obtained from ABclonal Technology. Antibodies against CD133, p44/42 MAPK (Erk1/2), phosphor-p44/42 MAPK (p-Erk1/2), and phosphor-Akt (p-Akt) were purchased from Cell Signaling Biotechnology. MYC antibody was obtained from Santa Cruz Technology. Mouse anti-GAPDH antibody was obtained from Sigma Aldrich. Immunoblot signals were acquired by the Amersham Imager 600 (General Electric Company).

### Image flow

SEM cells labeled with Cell Trace Far Red and CD19 CAR-Jurkat T cells labeled with copGFP were co-cultured at the ratio of 1:1 for 1 h. Cells were re-suspended in 4% PFA for 30 min, and images were acquired on the Amnis Imagestream Mk II Imagine flow cytometer (Luminex).

### Super-resolution imaging

REH cells expressing CD19-mRuby2 fusion and CD19 CAR-Jurkat T cells expressing copGFP were seeded into cell culture imaging dishes. Protease inhibitor cocktail was added to prevent CD19 antigen degradation. Images were acquired on the GE Delta Vision OMX SR imaging system. ImageJ software was used to generate the figures.

### qRT-PCR

qRT-PCR was performed using 7500 Real-Time PCR Systems (Applied Biosystems). The data represent relative mRNA levels normalized to *GAPDH*. Primers used for qRT-PCR assays were listed below:

CD19-Forward: CCCAAGGGGCCTAAGTCATTG,
CD19-Reverse: AACAGACCCGTCTCCATTACC;
GAPDH-Forward: GGCACAGTCAAGGCTGAGAATG,
GAPDH-Reverse: ATGGTGGTGAAGACGCCAGTA.

### Mouse studies and in vivo imaging

SEM cells simultaneously expressing GFP and luciferase were described previously.^10^ NOD/SCID mice were purchased from Vital River Laboratories. 1.5 million luciferase-expressing cells were intravenously injected via tail vein into NOD/SCID mice. NOD/SCID mice were then administered 1.5 million scFv-expressing MSCs on a twice-weekly schedule beginning 3 days after xenograft. Total body bioluminescence was quantified at indicated time points. All animal work was performed in accordance with a protocol approved by the Animal Studies Committee of Ruijin Hospital.

### Statistical analysis

All statistical analyses were performed using the GraphPad Prism 6 software. The Student’s *t*-tests were used to analyze the differences between the groups.

## Results

### Target cell death through a T-cell independent mechanism

Since cytotoxic CTLs or CAR-T cells mediated target cell destruction is an acute process, the in vitro capability of CTL or CAR-T cells is usually tested using a short time-specific cell lysis assay. Whether these effector cells have any chronic effects on target cells when co-cultured for an extended length of time has received scant attention to date.

In a co-culture system, B cell acute lymphoblastic leukemia (B-ALL) target cells caused CD19-CAR-Jurkat T cells to activate robustly and quickly, and the extent of T-cell activation was substantially reduced after 24 hours as demonstrated by an attenuation in ERK phosphorylation (Fig. 1A and S1A). However, despite the diminished activation of effector cells, apoptosis in target cells continuously increased after 24 hours (Fig. 1B). To address whether the sustainable death in the late time point was a result of the transient activation of T cells at an earlier time point, we collected target cells and effector cells separately after co-cultured for 24 hours, and then cultured with fresh effector cells or target cells, respectively. Interestingly, apoptosis continuously increased in the pre-co-cultured target cells when they were re-co-cultured with fresh CD19-CAR-Jurkat T cells, even though they could no longer effectively activate these effector cells (Fig. 1C, D). Importantly, the pre-co-cultured target cells showed only minimal apoptosis when they were re-co-cultured with fresh control Jurkat T cells, thus excluding the possibility that the T-cell activation mediated cytolytic activity seen at earlier periods were sufficient to cause the sustainable increase in apoptosis at the later period. These results suggested that target cell death, especially in the later period, might not be directly associated with T-cell activation. Interestingly, the pre-co-cultured CD19 CAR-Jurkat T cells could still be activated by freshly-added target cells (Fig. 1E), suggesting that the diminished activation of effector cells in the later period was caused by that the pre-co-cultured target cells became unable to activate the effector cells, rather than that the pre-co-cultured effector cells themselves became non-responsive to target cells.

**Figure 1.**
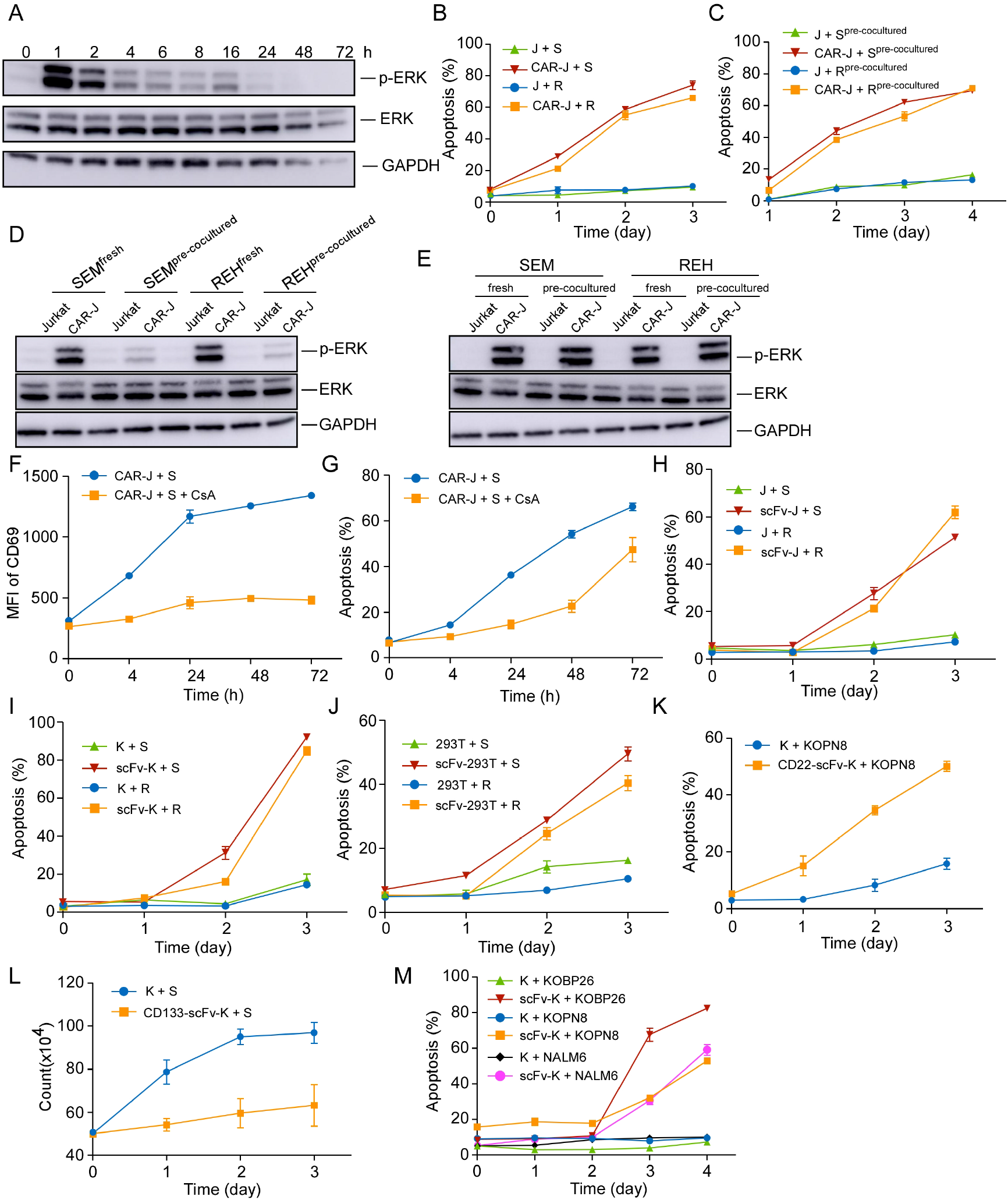
Target cell death through a T-cell independent mechanism. (A) The phosphorylated ERK levels of CD19 CAR-transduced Jurkat T (CAR-J) cells were determined at indicated times after co-cultured with SEM cells. (B) The specific lysis of CellTrace PE-labeled target cells was assayed by Annexin V staining at the indicated times. J, Jurkat; S, SEM; R, REH. (C) The effector cells were co-cultured with the target cells, and were then separately collected by FACS sorting (referred as pre-cocultured). The pre-cocultured target cells were re-cocultured with fresh effector cells and the specific lysis of the target cells were analyzed by Annexin V staining. (D) CD19-CAR-Jurkat or control Jurkat cells were co-cultured with fresh or pre-cocultured target cells for 1 h, and the phosphorylated ERK levels in the effector cells were evaluated by immunoblots. (E) The fresh or pre-cocultured CAR-J or control Jurkat T cells were co-cultured with the indicated fresh target cells for 1 h, and immunoblots of phosphorylated ERK levels in the effector cells were evaluated by immunoblots. (F) CD19-CAR-J cells were co-cultured with SEM cells with or without 200 nM CsA, and the mean fluorescence intensity (MFI) of the activation marker CD69 was evaluated by flow cytometry. (G) CD19-CAR-J cells were co-cultured with SEM cells with or without 200 nM CsA, and Annexin V staining of SEM cells at indicated time was detected. (H-J) CD19-scFv-expressing Jurkat (scFv-J), CD19-scFv-K562 (scFv-K), or CD19-scFv-293 (scFv-293) cells were co-cultured with CellTrace Far Red-labeled SEM or REH cells at the E/T ratio of 1:1. The apoptosis of the target cells was analyzed by Annexin V staining. scFv-transduced cells denote CAR-T cells that lacks co-stimulatory and CD3 zeta-chain signalling domains. (K) Annexin V staining of Kopn8 cells after co-cultured with CD22-scFv-K562 or control K562 cells at the ratio of 1:1. (L) CD133-scFv-K562 or control K562 cells were co-cultured with SEM cells at the ratio of 1:1. Cell count of SEM cells was examined at indicated times. (M) Apoptosis of indicated target cells after co-cultured with CD19-scFv-K562 or control K562 cells. Error bars reflect ± SEM.

To understand whether T-cell activation is absolutely required for CAR-T-cell-mediated target cell death, we suppressed the activation of CAR-T cells using the calcineurin inhibitor cyclosporin (CsA) and tested whether target cells could still be destroyed by these anergic CAR-T cells. As expected, while the activation of CAR-T cells was continuously blocked by CsA, as evidenced by diminished CD69 expression (Fig. 1F), a large portion of B-ALL cells still died when co-cultured for an extended time (Fig. 1G). To further determine whether the activation of T cells is necessary for target cell death, we determined whether target cells could be killed by non-signaling CD19-CAR-Jurkat (hereinafter referred as CD19-scFv-Jurkat) cells (Fig. S1B). Indeed, the B-ALL target cells could still undergo significant apoptosis when co-cultured with these CD19-scFv-Jurkat cells, albeit the death kinetics were much slower than that of CD19-CAR-Jurkat cells (Fig. 1B, H). These results suggest that there are mechanisms other than T-cell activation involved in the process of CD19-CAR-T mediated destruction of B-ALL cells.

Taken together, these results suggested that CAR-T cells, at least for CD19-CAR-T cells, can achieve target cell death through two distinct pathways, one involving classical CTL pathway, which causes rapid death and is dependent on T-cell activation, and the other, being of slower kinetics and is T-cell activation independent.

### Target cells can be destroyed by various artificial antigen-recognizing cells

Since the activation of T cells is not crucial for target cell death, we reasoned that this targeted cell death effect could be alternatively achieved by other non-lymphocyte-derived antigen-recognizing cells. To this end, we determined whether non-T-cell-derived CD19-scFv bearing effector cells could cause target cell death (Fig. S1B). Interestingly, the B-ALL targeted cells could also be killed by co-cultured CD19-scFv-K562 cells and CD19-scFv-293T cells, with similar death kinetic to CD19-scFv-Jurkat cells (Fig. 1I, J). These results suggested both lymphoid cells and non-lymphoid cells can induce efficient target cell death when they are armed with an artificial antigen recognizing module.

To further understand whether this is a general phenomenon, we tested the targeted killing capabilities for scFv-bearing K562 cells recognizing antigens other than CD19. Among these scFv-bearing K562 cells, CD22 targeting K562 cells could cause efficient target cell death in CD22-positive KOPN8 cells (Fig. 1K). While CD133 targeting K562 cells could only retard cell proliferation without causing efficient target cell death (Fig. 1L and S1C). Nevertheless, besides REH and SEM cells, other B-ALL cells such as KOPN8, KOBP26, and Nalm6 cells could also be effectively killed by CD19-scFv-K562 cells, suggesting that B-ALL cells can be generally destroyed by CD19-scFv-K562 cells (Fig. 1M). However, some CD19 positive B-cell malignancy cells such as RAJI cells could not be killed by these CD19-scFv-bearing effector cells (Fig. S1D-F). These results suggested that the ability of scFv-bearing effector cells to kill target cells is dependent on the target cell type and the target antigen selected. Together, these results suggested that it is feasible to utilize non-lymphocyte-derived artificial antigen-recognizing cells (AARCs) as effector cells to destroy or inhibit target cancer cells.

### Artificial antigen-recognizing cells cause antigen depletion on target cells

We next addressed the question of how AARCs can kill or inhibit target cancer cells. Since target cells can no longer activate effector CAR-T cells in the late period of co-culture, we suspected that the death or inhibition of target cells might be related to the antigen loss on target cells.

Previous studies have found that CD19-CAR-T cells provoke reversible CD19 antigen loss through trogocytosis, an active process in which the target antigen is transferred to T cells.^11^ We first monitored the interaction between effector cells and target cells, and surmised that CD19-CAR-Jurkat T cells and SEM cells could form conjugates in various formats (Fig. 2A and S2A). Since trogocytosis occurs at the immunological synapse and conjugate formation is a perquisite for trogocytosis,^12^ these conjugates could represent those cells undergoing trogocytosis. When SEM cells were co-cultured with CD19-CAR-Jurkat T cells, the cell surface CD19 was remarkably reduced on the conjugated SEM cells (Fig. 2B, C), which coincided with a slight increase of CD19 on CD19-CAR-Jurkat T cells (Fig. 2D), indicating that these cells did indeed undergo trogocytosis. To our surprise, the cell surface CD19 was also remarkably reduced on the unconjugated SEM cells, even when co-cultured at a low E/T ratio of 1:10 (Fig. 2B, C). To confirm that the diminished CD19 antibody labeling was a result of CD19 depletion, rather than a competition between CAR molecules and CD19 antibody,^13^ we examined whether CD19 was degraded using immunoblotting. Indeed, CD19 protein levels were significantly reduced in B-ALL target cells after co-cultured with CD19-CAR-J effector cells (Fig. 2E). Notably, the increase in CD19 on effector cells was merely transient and incomparable to those lost on target cells (Fig. 2B, D, F). These results suggested that the CD19 loss might not be principally through trogocytosis. To further exclude the possibility that these unconjugated SEM cells were actually those cells post-trogocytosis, SEM cells were co-cultured with non-signaling CD19-scFv-Jurkat T cells, which were incapable of mediating trogocytosis, since T-cell-mediated trogocytosis requires signaling of T cells.^12^ Interestingly, CD19 was consistently lowered on the target SEM cells (Fig. 2G). Moreover, CD19 depletion was also observed when co-cultured with CD19-scFv expressing K562 cells (Fig. 2H) or 293T cells (Fig. 2I). These results suggested that besides CAR-T cells, other artificial CD19-recognizing cells could also intrinsically cause cell surface CD19 depletion on the target cells through an unknown mechanism independent of lymphocyte-mediated trogocytosis.

**Figure 2.**
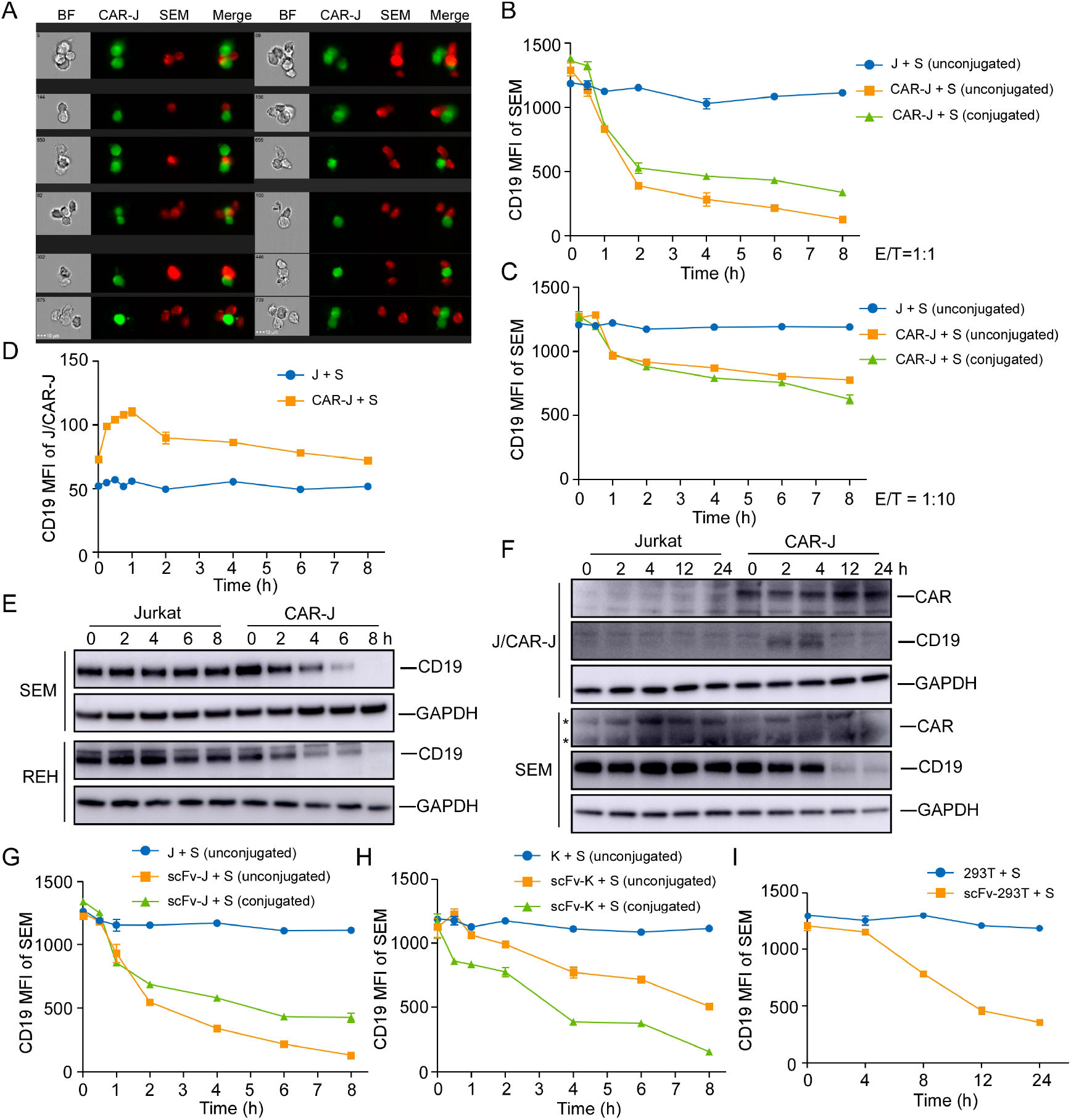
Artificial antigen-recognizing cells cause antigen depletion on target cells. (A) SEM cells were co-cultured with CD19-CAR-J cells for 1 h. Cells were monitored by image flow cytometry. BF, bright field. Scale bar, 10 μm. (B) MFI of CD19 in conjugated or unconjugated SEM cells after co-cultured with Jurkat or CAR-J cells. The conjugated and unconjugated cells were distinguished using FlowJo software. (C) MFI of CD19 in SEM cells co-cultured with Jurkat or CAR-J cells at the E:T ratio of 1:10. (D) MFI of CD19 in unconjugated Jurkat or CAR-J cells. (E) Immunoblots of target cells after co-cultured with Jurkat or CAR-J cells. (F) Immunoblots of SEM or Jurkat/CAR-J cells after co-cultured and sorted by flow cytometry. *, nonspecific bands. (G-I) MFI of CD19 in the indicated SEM cells following co-cultured with CD19-scFv-J (G), CD19-scFv-K (H) or scFv293T (I) cells. Error bars reflect ± SEM.

We next examined whether the phenomenon of antigen depletion could extend to other cell types and antigens. Even though CD19-scFv-K562 cells couldn’t kill RAJI cells, they could induce effective CD19 antigen depletion in RAJI cells (Fig. S2B). Furthermore, CD22 and CD133 targeting effector cells could also induce effective antigen depletion in the corresponding target cells (Fig. S2C-F). Thus, antigen depletion is a general feature of artificial antigen recognizing cells, which can probably be applied to many, if not all, antigens and cell types.

### CD19 depletion is mediated by endocytosis in target cells

Although CD19 became depleted upon exposure to CD19-targeting effector cells, there was little variation in the expression of *CD19* mRNA (Fig. 3A). Moreover, we found that diminished CD19 expression in the pre-co-cultured SEM cells was reversed without the presence of CD19-targeting effector cells (Fig. 3B), indicating a reversible and post-transcriptional mechanism was responsible for the CD19 depletion caused by the presence of artificial CD19-recognizing cells. Interestingly, the status of CD19 depletion in pre-co-cultured SEM cells was maintained in the presence of CD19-CAR-Jurkat effector cells (Fig. 3B), even though they could no longer effectively activate these effector cells (Fig. 1D), further supporting our view that CD19 depletion caused by the presence of artificial CD19-recognizing cells is T-cell activation independent.

**Figure 3.**
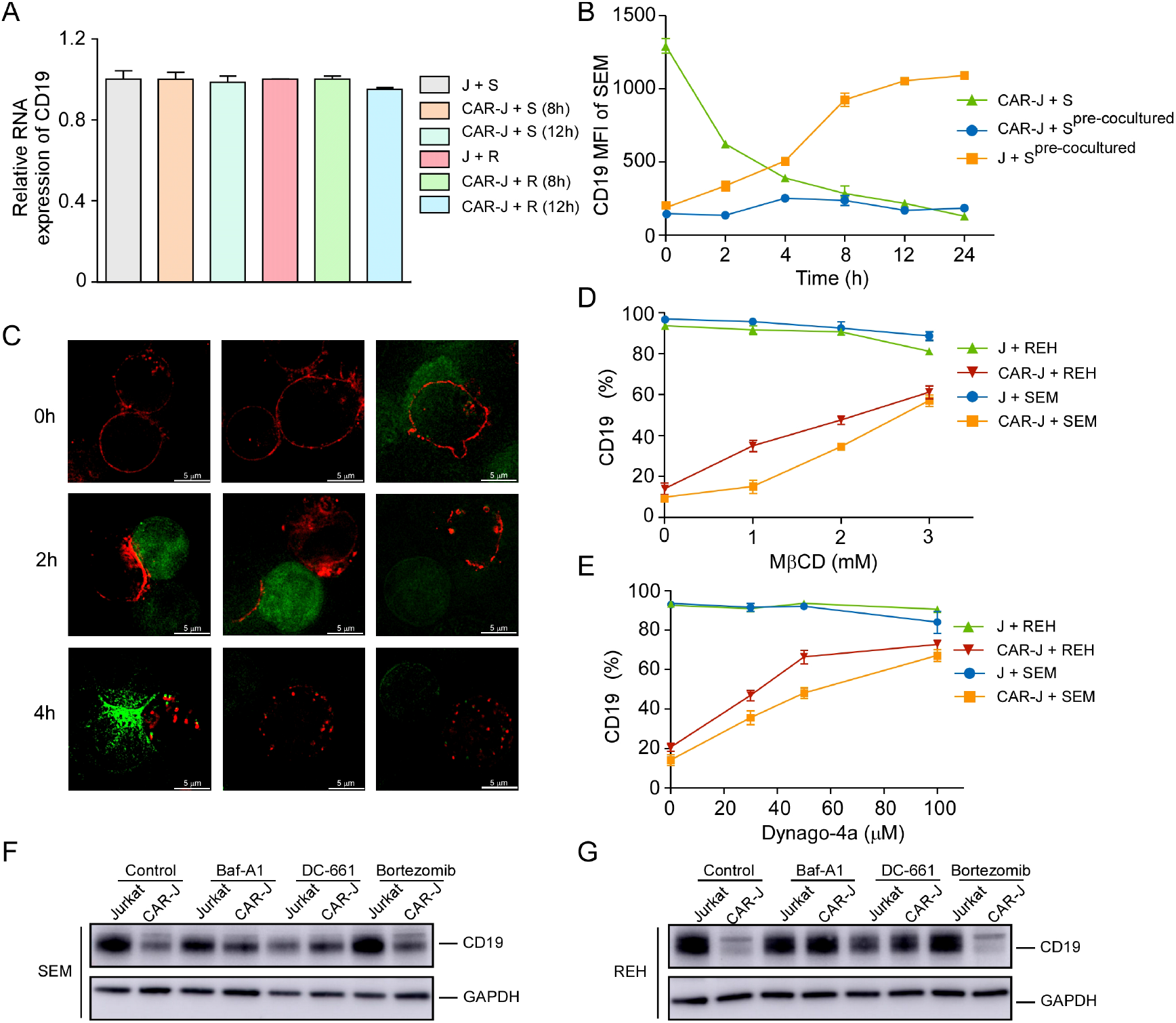
CD19 depletion is mediated by endocytosis in target cells. (A) SEM or REH cells were co-cultured with CAR-J or control Jurkat cells for the indicated time. The relative *CD19* mRNA levels in the target cells were evaluated by qRT-PCR. Values were normalized against *GAPDH*. (B) SEM cells were co-cultured with CAR-J cells for 8 h and then the target cells were sorted by flow cytometry. Fresh SEM cells or pre-cocultured SEM cells were re-cocultured with CAR-J or control Jurkat cells and the expression of CD19 in the target cells was analyzed by flow cytometry. (C) SEM cells were co-cultured with CAR-J cells and the fate of CD19 protein was monitored by DeltavVison OMX SR imaging system. The protease inhibitor cocktail was added to prevent protein degradation. Red color denotes CD19-mRuby2; Green color indicates CAR-J cells. (D, E) REH or SEM cells were co-cultured with indicated effector cells with exposure to MβCD (16 h) or dynago-4a (24h). The percentage of CD19 positive cells was detected. (F, G) SEM or REH cells were pre-treated with Baf-A1 (100 nM), DC-661 (10 μM) or Velcade (10 nM) for 2 h, then co-cultred with Jurkat or CAR-J cells. The levels of CD19 in the SEM (F) or REH (G) cells was analyzed by immunoblots. Error bars reflect ± SEM.

To address the detailed mechanism of CD19 depletion, we first tracked the behavior of CD19 proteins in the co-culture system of SEM and CD19-CAR-Jurkat cells. To this end, CD19-mRuby2 fusion proteins were ectopically expressed in REH cells (Fig. S3A), and their degradation was prevented by the protease inhibitor cocktail. In control target cells, CD19 proteins were found to be distributed evenly on the cell surface (Fig. 3C). After co-cultured with effector cells for 2 hours, however, CD19 in the target cells tended to coalesce to discrete cell surface areas which were enriched at the interface with effector cells (Fig. 3C). Interestingly, CD19 proteins in both conjugated and unconjugated target cells showed a similar pattern, indicating that a transient interaction with effector cells was sufficient to cause this translocation of CD19 protein. After an additional 2 hours, CD19 proteins translocated further into discrete foci inside the target cells (Fig. 3C), suggesting that they were internalized into the target cells. To determine whether the internalization of CD19 was mediated by endocytosis, we examined whether CD19 depletion on the cell surface of target cells could be prevented by various inhibitors of endocytosis including cholesterol depleting methyl-β-cyclodextrin (MβCD)^14^ and the dynamin inhibitor Dyngo-4a. Indeed, CD19 internalization and depletion could be rescued by these endocytosis inhibitors in a dose-dependent manner (Fig. 3D, E), confirming that the internalization of CD19 is mainly mediated by endocytosis.

The depletion of CD19 after co-culture suggested that this endocytosis of CD19 was ultimately terminated. One of the main degradation pathways for endocytosed proteins is lysosomal degradation.^15^ Therefore, to evaluate the pathway causing CD19 degradation after endocytosis, we examined the effects of various lysosomal and proteasomal inhibitors on CD19 depletion. We found that the lysosomal inhibitors Bafilomycin A1 and DDC-61 significantly abolished the CD19 depletion caused by co-culture with CD19-CAR-J cells, while the proteasome inhibitor bortezomib had little effect (Fig. 3F, G). Collectively, our results suggested that the CD19 antigen is internalized and degraded by lysosomes inside the target cells in the presence of CD19-targeting effector cells.

### CD19 depletion undermines the survival of B-ALL target cells via sabotage of the CD19/AKT/MYC axis

As a cell surface signaling protein, CD19 is required for several processes involved in B-cell development and function, and by inference, is important for the survival of malignant B cells, and serves as a crucial therapeutic target for B-cell malignancies.^16–18^ Since the death of CD19 positive B-ALL cells caused by CD19-targeting AARC, is accompanied by the depletion of CD19 in the target cells, we reasoned that CD19 depletion might account for the T-cell-activation independent chronic death effect of CD19-AARC. Indeed, acute CD19 depletion using CD19-targeting shRNAs led to apoptosis in REH and SEM cells (Fig. 4A and S4A), suggesting competent CD19 signaling is indispensable for the survival of these B-ALL cells, and CD19 depletion caused by CD19-AARC cells may lead to target cell death. In contrast, acute CD19 depletion couldn’t kill RAJI cells (Fig. S4B, C), suggesting competent CD19 signaling is dispensable for the survival of RAJI cells. These results might explain why CD19-scFv bearing AARC cells could cause CD19 depletion but not cell death in RAJI cells.

**Figure 4.**
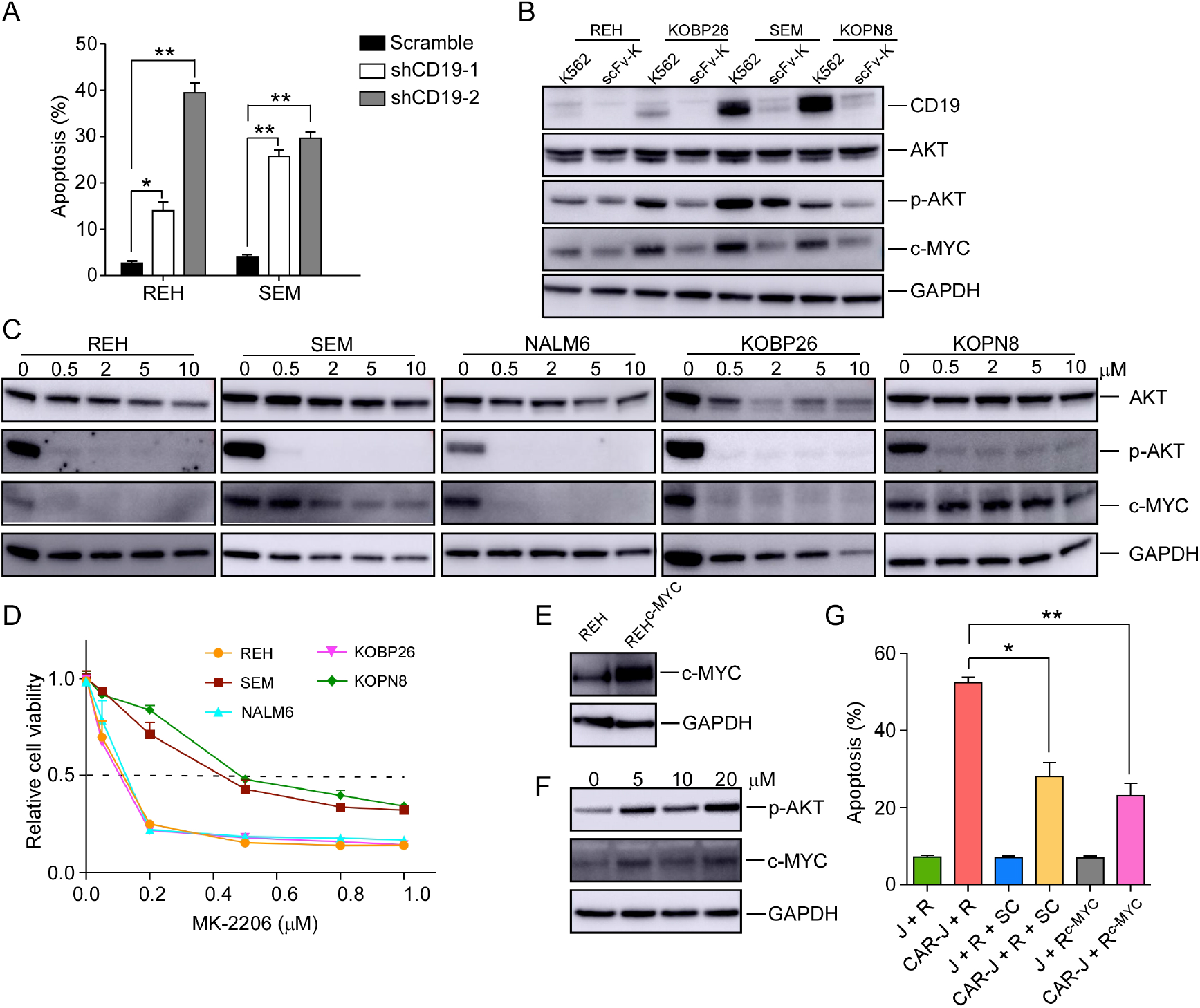
CD19 depletion undermines the survival of B-ALL target cells via sabotage of the CD19/AKT/MYC axis. (A) Apoptosis analysis of REH or SEM cells infected with the indicated lentiviral vectors for 48 h. (B) Immunoblots of target cells co-cultured with CD19-scFv-K562 or control K562 cells. (C) Immunoblots of the indicated cells treated with the Akt inhibitor MK-2206 for 24 h. (D) Cell viability of the indicated cells treated with MK-2206 at the indicated concentrations for 5 days. (E) Immunoblots of REH cells infected with the indicated lentiviral vectors. (F) Immunoblots of the indicated cells treated with the Akt activator sc-79. (G) Jurkat or CAR-J cells were co-cultured with the indicated target cells (R, REH; R+SC, REH cells treated with sc-79; R^c-Myc^, c-Myc overexpressed-REH cells). Apoptosis analysis of the indicated REH cells was detected by flow cytometry. Statistical analysis was performed using two-tailed *t*-test. Error bars reflect ± SEM.

The PI3K/AKT pathway is instrumental in BCR signaling in B cells. High expression of CD19 on the surface of malignant B cells may activate AKT, and up-regulate MYC to support the survival of these cells.^18, 19^ MYC is one of the key oncoproteins in B cell malignancies, and MYC dysfunction appears to uniformly reduce cell proliferation and induce apoptosis.^20^ We determined whether the PI3K/AKT pathway and MYC expression in the target cells may be compromised by CD19-AARC-mediated CD19 depletion. To this end, we examined the level of phospho-AKT and MYC in the B-ALL target cells after co-cultured with CD19-scFv-K562 cells. As expected, decreased MYC levels were accompanied by the reduction of phospho-AKT in CD19-depleted B-ALL cells (Fig. 4B). To confirm whether the dysfunction in the AKT/MYC axis could cause cell death in B-ALL cells, we treated the cells with the AKT inhibitor MK-2206. With the deactivation of AKT, MYC protein levels were accordingly reduced in REH, Nalm6 and Kobp26 cells, which correlates with significantly reduced cell survival, and confirms that MYC down-regulation could cause cell death in these cells (Fig. 4C, D). Interestingly, in SEM and KOPN8 cells whose survival has been shown to be dependent on MYC,^21^ MYC protein levels were not accordingly reduced with AKT deactivation. Instead, MYC protein levels were reduced only when high concentration of MK-2206 was used, which correlates with relative resistance to MK-2206, suggesting that the CD19 depletion mediated down-regulation of MYC levels in these cells might also engage other unknown AKT-independent pathways which can be inhibited by high concentration of MK-2206 (Fig. 4B-D). To further confirm that MYC downregulation is essential for B-ALL target cell death, we examined whether the reestablishment of MYC protein levels could rescue the target cell death. As expected, overexpression of MYC or rescuing the AKT/MYC axis using the AKT activator SC-79 in REH cells could mitigate the target cell death by CD19-scFv-K562 in target cells (Fig. 4E-G). These results therefore have established that AARC-mediated CD19 depletion and MYC destruction can cause cell death of target cells, which is mainly through disruption of the CD19/AKT/MYC axis.

### In vivo efficacy of CD19-AARC in a B-ALL xenograft mouse model

We next determined the *in vivo* efficacy of CD19-AARC cells in a previously established B-ALL xenograft mouse model.^22^ To this end, mesenchymal stromal cells (MSCs) were chosen as the source of AARC because of their low immunogenicity, widely demonstrated clinical safety, good transplantability, and tumor-homing features.^23^ We examined the *in vitro* and *in vivo* efficacies of CD19-scFv presenting MSCs. Similar to other CD19-AARC cells, CD19-scFv-MSCs caused CD19 depletion efficiently, as well as apoptosis in the co-cultured B-ALL target cells (Fig. 5A-C). Subsequently, mice transplanted with SEM cells were treated with CD19-scFv-MSCs at the indicated interval and leukemia progression was monitored using bioluminescence (Fig. 5D). Leukemic cells in the engrafted mice expanded rapidly in both untreated and control MSC treated conditions. In contrast, the engrafted mice treated with CD19-scFv-MSCs showed a striking reduction in leukemia progression (Fig. 5E, F). Our results demonstrated both the *in vitro* and *in vivo* efficacies of CD19-scFv-MSCs, suggests that CD19-AARC cells can serve as a novel cellular therapeutic approach in the treatment of B-ALL leukemia.

**Figure 5.**
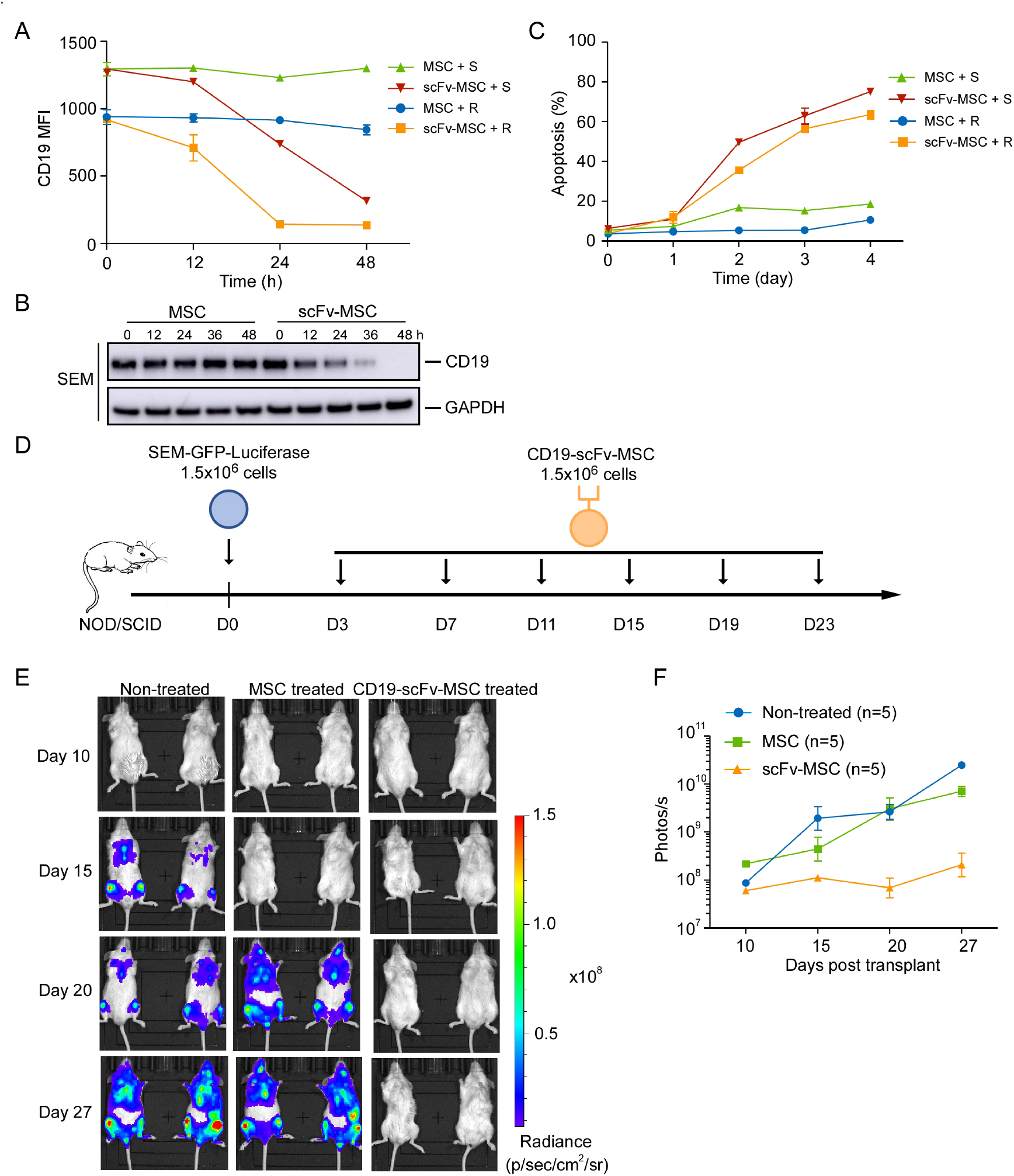
In vivo efficacy of CD19-AARC in a B-ALL xenograft mouse model. (A) REH or SEM cells were co-cultured with CD19-scFv-transduced MSCs (CD19-scFv-MSC) or control MSC cells at the indicated times, and the levels of CD19 in the target cells were determined by flow cytometry. (B) Immunoblots of SEM cells co-cultured with CD19-scFv-MSC or control MSC cells for the indicated time. (C) The apoptosis of SEM or REH cells cocultured with CD19-scFv-MSC cells was determined by Annexin V staining. (D) Treatment schedule for administration of CD19-scFv-MSCs. (E) NOD/SCID mice transplanted with luciferase-expressing SEM cells were treated with CD19-scFv-MSCs. On the indicated days after xenograft, mice were imaged to assess for leukemia progression. Representative bioluminenscence images are shown on the left and the quantification of bioluminescence (photonic flux) over the duration of treatment is shown on the right. Error bars reflect ± SEM.

## Discussion

Targeted cell death by CTLs is thought to be a rapid process. Therefore, a 4-hour cytotoxicity assay is routinely used to test the *in vitro* efficacy of CAR-T cells. However, the chronic effect of CAR-T cells has seldom been addressed. Here we have shown that the cytotoxic effect of CAR-T cells is composed of both acute and chronic effects (Fig. 6). The former is rapid and dependent upon T-cell activation, while the latter has much slower kinetics and is independent of T-cell activation but instead relies on the depletion of the corresponding antigen. Importantly, only a fraction of the target cells may form stable associations with effector cells to initiate cell death through the classical acute CTL pathway. In contrast, most of the target cells can be deprived of the corresponding antigens through a transient interaction with effector cells and be destroyed through the chronic pathway. Thus, ideal CAR-T therapeutics with long-term efficacy should have a good chronic effect.

**Figure 6.**
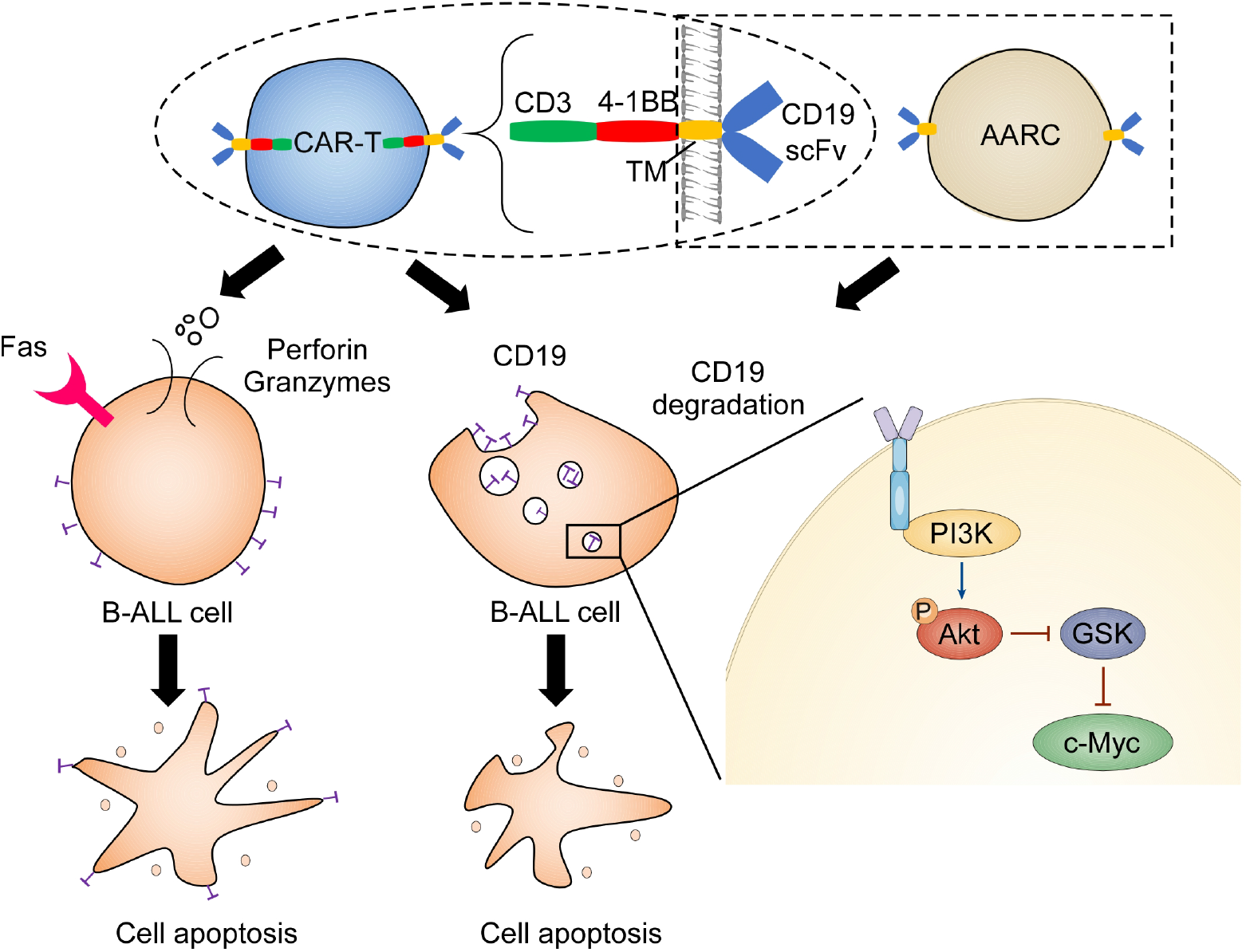
Illustration depicts how CAR-T cells induce CD19 degradation and CD19-dependent cell death.

Since the chronic effect of CAR-T cells is largely dependent on the antigen selected, our study thus provides valuable insight into antigen selection and CAR design. The first choice is that the antigen won’t be internalized and depleted when recognized by the corresponding CAR. In this way, the escape of the antigen can be avoided and the target cells can be continuously recognized and destroyed. Alternatively, the antigen should be critical for the survival of the target cells, and can be internalized and depleted when being recognized by effector cells. Thus, dysfunction of the antigen can lead to cell destruction. Here we showed CD19 is a superior target because CD19-recognizing effector cells can maintain the target cells being deprived of cell-surface CD19, and therefore effectively cause chronic cell death in the target cells whose survival is dependent on CD19 signaling. This can partly explain why CD19-CAR-T remains the most successful CAR-T therapeutics. Since various antigens may have differential characteristics in different cancer cells, this at least in part explains why different cancers respond differently to different CAR-Ts, or even the same CAR-T.

Together with previous reports,^11^ our results certainly suggest for some antigens such as CD19, that antigen escape is inevitable. However, according to the dependency of the cells to antigens, this antigen escape can also be manipulated into a cell death approach, despite that it may also prevent the cells from becoming recognized by CAR-T cells. Although acute depletion of CD19 is detrimental to B-ALL cells, they can transform into an adaptive state, or clones with a growth advantage, so that their survival is no longer dependent on CD19.^16, 24^ Based on our mechanical findings, the resistant cells might be conferred with the ability to survive in the absence of CD19. This might partly explain why different B cell malignancies have different relapse rates after CD19-CAR-T therapy.^25^

The AARC could maintain the antigen depletion status of target cells through transient interaction, which can endow them with cell death capability as a serial killer. Although antibody-based drugs such as CD19 antibodies can also cause antigen depletion,^26^ and therefore, kill or inhibit the target cells through disrupting the function of the antigen,^18^ they can become nullified since antibodies would be co-internalized with their antigens. This implies that to achieve sustainable antigen depletion status, antibodies have to add in excess or constantly infused to neutralize the recovery of the antigen. Therefore, for this aspect of function, AARC as a living drug should transcend antibodies as a consumable drug.

## Acknowledgments

This work was supported by the National Key Research and Development Program of China (2018YFA0107802), the National Major Scientific and Technological Special Project (2018ZX09101001), the National Natural Science Foundation of China (81973996 and 81570119), the Program of Shanghai Academic Research Leader (19XD1402500), the Shanghai Municipal Education Commission Gaofeng Clinical Medicine Grant (20161304), the Shanghai Municipal Health Commission (2019CXJQ01), the Open Project Program of the National Facility for Translational Medicine (Shanghai) (TMSZ-2020-204), the Open Project Program of Ministry of Education Engineering Research Center of Cell & Therapeutic Antibody (19X110020009-002), the China Postdoctoral Science Foundation (2020M681339 and 2020M681338), the Collaborative Innovation Center of Hematology, and the Samuel Waxman Cancer Research Foundation.

## Authors’ contributions

D.L. and H.L. conceived the project and designed the experiments. D.L., R.H.W., W.B.W., M.L.G. and S.F.X., performed most of the experiments. M.L.G., Q.Y.X., and Y.S., performed some experiments and provided technical assistance for the mouse experiments. D.L., M.L.G., J.Z., and H.L. analyzed the data. D.L., M.L.G., J.Z., and H.L. wrote the paper. All authors discussed the results and commented on the manuscript.

## Footnotes

Any data and materials that can be shared by the corresponding author will be released freely.

