## Supplemental Figures for "A T-cell independent universal cellular therapy strategy through antigen depletion"

### Supplementary Figures

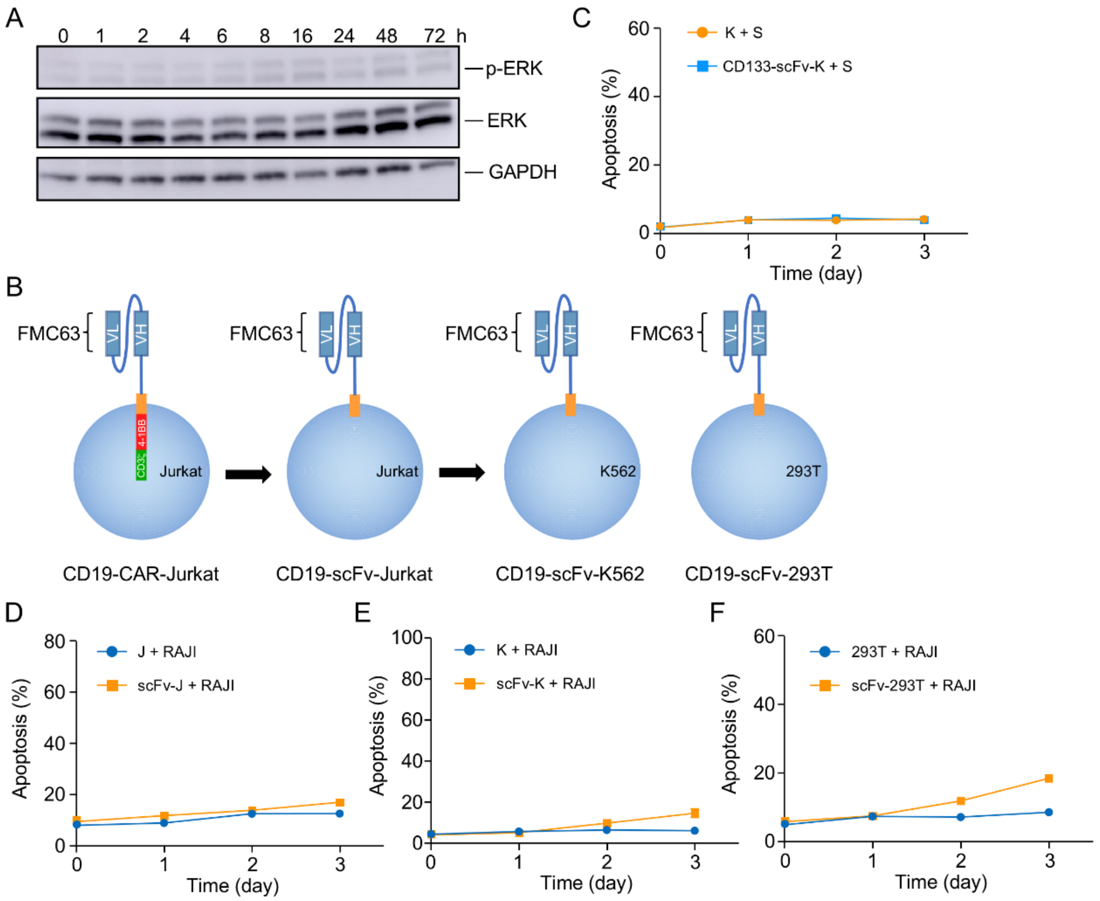


**Fig. S1 Target cell death through a T-cell independent mechanism.** (A) The phosphorylated ERK levels of Jurkat T cells were determined after co-cultured with SEM cells for the indicated times. (B) The construction sketches of different effector cells. (C) Annexin V staining of SEM cells after co-cultured with CD133 scFv-K562 cells at the ratio of 1:1. (D-F) CD19 scFv-expressing Jurkat (scFvJ), CD19 scFv-K562 (scFvK), or CD19 scFv-293 (scFv293) cells were co-cultured with CellTrace Far Red-labeled RAJI cells at the ratio of 1:1. The apoptosis of target cells was analyzed.

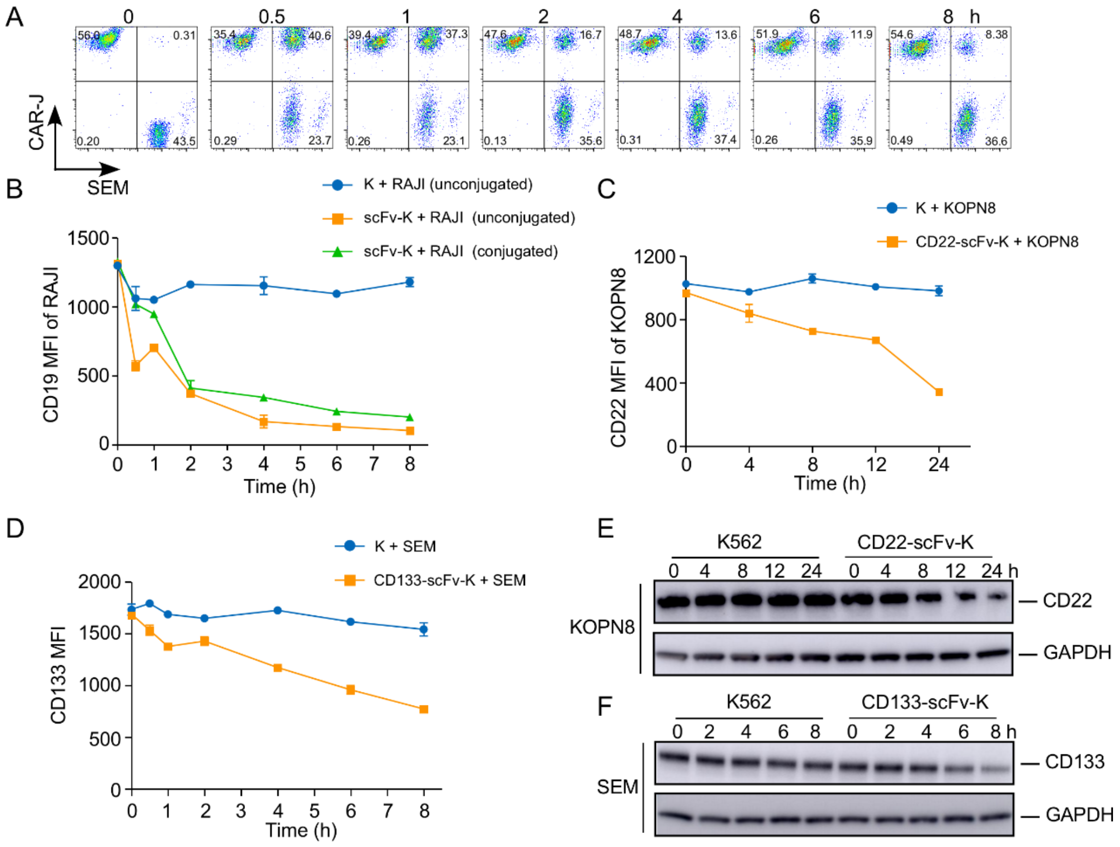


**Fig. S2 The phenomenon of antigen depletion on other antigens.** (A) SEM cells were co-cultured with CD19 CAR Jurkat T cells, and conjunction was detected at indicated times. (B) MFI of CD19 in RAJI cells was detedcted at indicated time. (C, D) CD22 scFv-K562 (CD22 scFvK), or CD133 scFv-K562 (CD133 scFvK) cells were co-cultured with the indicated target cells at the ratio of 1:1. The expression of CD22, and CD133 of target cells was analyzed. (E, F) Immunoblots of the indicated antibodies in target cells after co-cultured with the indicated effector cells.

A

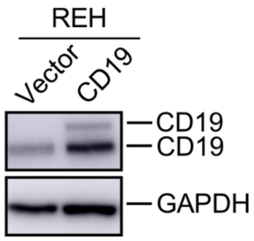


**Fig. S3 The expression of CD19.** (A) The CD19 protein levels of REH cells infected by the indicated lentiviral vectors.

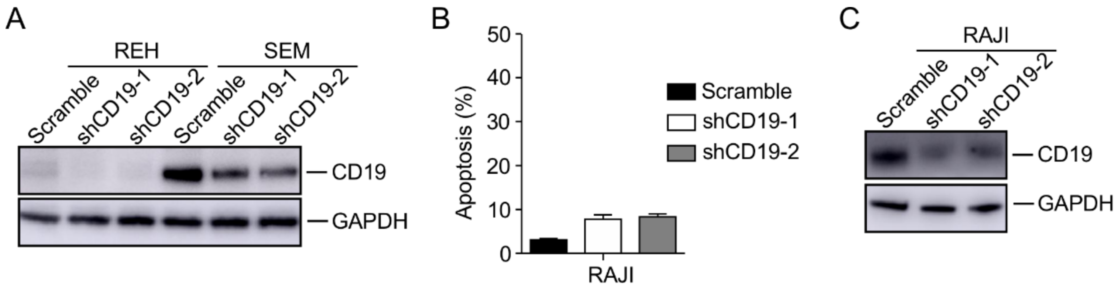


**Fig. S4** (A) The CD19 protein levels of REH and SEM cells infected by the indicated lentiviral vectors. (B) Apoptosis analysis of RAJI cells infected with the indicated lentiviral vectors for 48 h. (C) The CD19 protein levels of RAJI cells infected by the indicated lentiviral vectors.
